# Mitochondrial oxidant stress promotes α-synuclein aggregation and spreading in mice with mutated glucocerebrosidase

**DOI:** 10.1101/2024.06.13.598820

**Authors:** Pietro La Vitola, Eva M Szegö, Rita Pinto-Costa, Angela Rollar, Eugenia Harbachova, Anthony HV Schapira, Ayse Ulusoy, Donato A Di Monte

## Abstract

Mutations of the glucocerebrosidase-encoding gene, *GBA1*, are common risk factors for Parkinson’s disease. Although only a minority of mutation-carrying individuals develops the disease, the mechanisms of neuronal vulnerability predisposing to pathology conversion remain largely unclear. In this study, heterozygous expression of a common glucocerebrosidase variant, namely the L444P mutation, was found to exacerbate α-synuclein aggregation and spreading in a mouse model of Parkinson-like pathology targeting neurons of the medullary vagal system. These neurons are primary sites of α-synuclein lesions in Parkinson’s disease and were shown here to become more vulnerable to oxidative stress after L444P expression. Nitrative burden paralleled the enhanced formation of reactive oxygen species within vagal neurons expressing mutated glucocerebrosidase, as indicated by pronounced accumulation of nitrated α-synuclein. A causal relationship linked mutation-induced oxidative stress to enhanced α-synuclein pathology that could indeed be rescued by neuronal overexpression of the mitochondrial antioxidant enzyme superoxide dismutase 2. Further evidence supported a key involvement of mitochondria as sources of reactive oxygen species as well as targets of oxidative and nitrative damage within L444P-expressing neurons. Scavenging of oxygen species by superoxide dismutase 2 effectively counteracted deleterious nitrative reactions and prevented nitrated α-synuclein burden. Taken together, these findings support the conclusion that enhanced vulnerability to mitochondrial oxidative stress conferred by glucocerebrosidase mutations should be considered an important mechanism predisposing to Parkinson’s disease pathology, particularly in brain regions targeted by α-synuclein aggregation and involved in α-synuclein spreading.

## Introduction

*GBA1* is the gene encoding the lysosomal enzyme glucocerebrosidase (GCase) that catalyses the hydrolysis of glucosylceramide to glucose and ceramide. The critical role played by GCase in preventing substrate accumulation, avoiding lipid dysmetabolism and preserving cellular structures and functions is indicated by the deleterious consequences of homozygous *GBA1* mutations that cause Gaucher disease, the most common lysosomal storage disease^1^. Genetic and clinical investigations over the past two decades have also established an intriguing relationship between *GBA1* mutations and Parkinson’s disease (PD)^2–4^. Five-10% of patients with idiopathic PD, rising to 25% in Ashkenazi patients, are found to carry a *GBA1* mutation/variant, underscoring not only the frequency of *GBA1*-associated PD but also the clinical similarities between this form of parkinsonism and the idiopathic disease^3–5^. *GBA1* mutations act as archetypal risk factors since only a percentage (10 to 30%) of individuals carrying these mutations ultimately develop PD^6,7^. Mechanisms underlying disease pathogenesis in affected mutation carriers remain largely unclear, and this knowledge gap significantly limits our ability to develop predictive markers of pathology conversion and potential strategies for early therapeutic intervention.

*GBA1* mutations are also risk factors for another progressive brain disorder, namely dementia with Lewy bodies that, similar to PD, is characterised by the accumulation of α-synuclein (ASYN)-containing neuronal inclusions (called Lewy bodies and Lewy neurites) and is therefore considered a synucleinopathy^8^. This observation supports the notion that formation of ASYN inclusions represents a common phenotype of pathological processes triggered by *GBA1* mutations. It also suggests that changes in ASYN expression, metabolism and/or toxic properties may be key events linking *GBA1* mutations and loss of GCase activity to the development of human synucleinopathies. Two toxic properties of ASYN, namely its ability to self-assemble into fibrillar aggregates and its inter-neuronal mobility, may explain features and mechanisms of ASYN pathology. Protein aggregation is likely to underlie the formation of ASYN-containing Lewy inclusions, while neuron-to-neuron ASYN transmission could contribute to the progressive spreading of pathology in PD and other synucleinopathies^9,10^.

Investigations into the relationship between *GBA1* mutations and ASYN aggregation and spreading have yielded significant clues but also raised important questions. Loss of GCase activity has been reported to promote ASYN misfolding and assembly, possibly due to a relationship between GCase reduction, increased intracellular ASYN levels and abnormal ASYN-lipid interactions^11–15^. A direct link between GCase deficiency and ASYN aggregation is not consistent with the findings of other investigations, however. For example, chemically induced enzyme depletion failed to produce any evidence of ASYN aggregation in primary hippocampal neuron cultures^16^. Furthermore, in mice with heterozygous expression of the L444P mutation of the *Gba1* gene (L444P/wild-type, L444P/wt, mice), a 30-40% decrease in GCase activity was not associated with pathological α-synuclein assembly, nor did it cause other PD-like abnormalities^17–19^. Taken together, these findings suggest that, even in experimental models of ASYN pathology, GCase mutations act as risk factors that predispose to but do not necessarily cause ASYN aggregation. Aggregate pathology would therefore require the contribution of other mutation-associated mechanisms that remain relatively unclear.

The relationship between PD-linked GCase mutations and ASYN spreading has been assessed by a relatively small number of *in vivo* studies. In these studies, induction of ASYN pathology and its progressive brain propagation were achieved by intraparenchymal administration of pathogenic pre-formed ASYN fibrils (PFFs)^20–23^. PFFs were injected, for example, unilaterally or bilaterally into the striatum of L444P/wt mice or unilaterally into the olfactory bulb of mice carrying a heterozygous D409V mutation of *Gba1*. PFF-induced pathology was reported to be enhanced either in several brain regions or only in the hippocampus of L444P/wt mice; it was instead unchanged or even reduced in the brain of D409V transgenic animals^20,21,23^. Variabilities in the results of these earlier studies may be due to differences in experimental tools (e.g., PFFs generated under non-identical conditions), procedures (e.g., single *vs.* double intraparenchymal injections) and/or transgene expression (L444P *vs.* D409V mutation). They nonetheless underscore the need for further investigations that may utilise different experimental paradigms of ASYN brain spreading and address unexplored questions, such as the nature of mutation-associated mechanisms capable of modulating inter-neuronal ASYN transfer.

Here, the effects of neuronal expression of the L444P GCase mutation were assessed using a well-characterized mouse model of ASYN pathology. In this model, protein aggregation and caudo-rostral brain spreading of ASYN are triggered by targeted overexpression of human ASYN (hASYN) within neurons of the dorsal medulla oblongata (dMO)^24–26^. First, ASYN pathology was compared in control *vs* L444P/wt. mice. Results of these experiments unequivocally indicated a marked exacerbation of both aggregation and spreading associated with L444P heterozygosity. A subsequent set of experiments was then aimed at elucidating mechanisms underlying this increased susceptibility to pathology. New evidence reported in this study revealed a key role of mitochondrial oxidative (Ox) and nitrative (Nt) stress in promoting mutation-induced ASYN aggregation and spreading. Rescue experiments were designed to counteract this stress and demonstrated that lowering the burden of Ox and Nt reactions effectively reversed the ASYN pathology triggered by neuronal L444P expression.

## Materials and methods

### Viral vectors

Adeno-associated viral vectors (AAVs) encoding for hASYN or superoxide dismutase 2 (SOD2) and empty-AAVs were generated using an AAV2-derived backbone plasmid (Supplementary Fig. 1) and AAV6-derived capsid. Expression of hASYN and SOD2 were driven by the promoter sequence for synapsin and CAG, respectively. A woodchuck-hepatitis virus post-transcriptional regulatory element (WPRE) and a polyadenylation signal sequence were inserted downstream to the promoter and protein encoding sequence. Empty-AAVs contained a synapsin promoter, WPRE and the polyadenylation signal sequence but lacked a protein coding sequence. AAV production and titration were performed by either Sirion Biotech (hASYN-AAVs) or Vector Biolabs (SOD2- and empty-AAVs). High-titer stock AAV preparations were diluted in phosphate-buffered saline to achieve a working titer of 3×10^11^ genome copies/ml (hASYN) or 2×10^12^ genome copies/ml (SOD2 and empty-AAVs).

### Animals and procedures

All animal protocols (81-02.04.2020.A169 and 81-02.04.2020.A277) were approved by the State Agency for Nature, Environment and Consumer Protection in North Rhine Westphalia, Germany. Heterozygous knock-in L444P mice (B6;129S4-Gbatm1Rlp/Mmnc) and non-transgenic littermates were obtained from the Mutant Mouse Regional Resource Centre (MMRRC_000117-UNC). L444P/wt mice were identified by PCR of the genomic DNA using the following forward and reverse primers: 5’-CCCCAGATGACTGATGCTGGA-3’ and 5’-CCAGGTCAGGATCACTGATGG-3’ (dx.doi.org/10.17504/protocols.io.5qpvo3xobv4o/v1). PCR amplification products were digested by the restriction enzyme *NciI*, yielding two fragments at 386 and 200 bp in heterozygous animals. Mice were kept in a specific pathogen-free animal facility on a 12 h light/dark cycle with ad libitum access to food and water.

Targeted overexpression of hASYN-, hASYN/empty- or hASYN/SOD2-AAVs in the dMO was induced in control and transgenic 9-to-14-month-old mice. A solution containing the AAV preparation was injected into the left vagus nerve as previously described^25,27^. Following surgery, mice were returned to their home cages and treated with pain killer medication for 3 days; they were then monitored daily throughout the duration of the experiments. No changes in body weight, basic motility, and general welfare were noticed as consequences of the surgical procedures. Six weeks after AAV administration, mice were sacrificed with an intraperitoneal injection of sodium pentobarbital (600 mg/kg). Brains were collected and either fresh-frozen for biochemical assays or perfused intracardially with 4% (w/v) paraformaldehyde for immunohistochemical observations. The perfused brains were immersed in 4% paraformaldehyde for 24 hours before cryopreservation in 30% (w/v) sucrose. Coronal sections of 35 μm were used to perform all histological analyses.

To visualize and assess intraneuronal reactive oxygen species (ROS) formation, subsets of AAV-treated control and transgenic mice received a subcutaneous injection of the superoxide indicator DHE immediately (1 hour) before the time of sacrifice^28^. Dihydroethidium (DHE) (Abcam, ab145360) was dissolved in 50% DMSO and injected at a dose of 10 mg/kg.

### GCase activity assay

The entire brain of control and transgenic mice was homogenized with a handheld homogenizer in lysis buffer (50 mM Tris-HCl, pH 7.4 and 750 mM NaCl, 5 mM EDTA and 10% Triton X-100). Homogenates were centrifuged at 5000 g for 10 min (4°C,), and protein concentration was assayed in the supernatants using the Pierce BCA Protein Assay Kit (Fisher Scientific, 23225). GCase activity measurements were performed as previously described^17^. Briefly, a standard curve was generated with serial dilutions of 4-methylumbelliferone, the product of enzymatic GCase activity measured in this assay. Samples were assayed in duplicates and quantified using a fluorescence spectrometer (365 nm excitation and 450 nm emission) (FLUOstar Omega, BMG LabTech).

### Western blot

Total (mouse and human) ASYN levels were measured using fresh frozen tissue collected from untreated and AAV-injected mice (dx.doi.org/10.17504/protocols.io.rm7vzjb22lx1/v1). The dMO was microdissected from tissue sections using a cryostat, and dissected samples were mechanically homogenized (Precellys 24 Touch Homogenizer, Bertin Technologies) with 0.5 mm Zirconium oxide beads in ice-cold lysis buffer; the buffer contained 1% Triton X-100 in phosphate-buffered saline solution (0.1 M, pH 7.6) supplemented with protease and phosphatase inhibitors. Samples were centrifuged (14000 g, 30 min, 4 °C), and 4 µg protein lysates (boiled for 5 min at 95 °C in Laemmli sample buffer supplemented with 5 % beta-mercaptoethanol) were loaded onto a 4-20% Tris/glycine SDS gel for western blot analysis. After blocking, membranes were incubated first in the presence of antibodies against ASYN (1:2000; BD Transduction Laboratories, RRID:AB_398108) and β-actin (1:10000; Abcam, RRID:AB_2305186), then with horse radish peroxidase-conjugated goat anti-mouse (ASYN) and goat anti-rabbit (β-actin) secondary antibodies (1:5000; Abcam, RRID:AB_955413 and RRID:AB_955417). The signal was visualized with a chemiluminescent substrate and detected with BioRad ChemiDoc. ImageJ software (ImageJ 1.54f, RID:SCR_003070, https://imagej.net/) was used to determine the optical density of protein bands (Gels plugin).

### Immunohistochemistry and spreading analysis

To assess caudo-rostral hASYN spreading, immunohistochemistry with brightfield detection was performed in serial coronal brain slices encompassing one-fifth of the whole brain^27^.. Briefly, free-floating sections were incubated in a quenching solution (3% H_2_O_2_ and 10% methanol in tris-buffered saline, pH 7.6). Non-specific binding was blocked by incubating samples in 5% normal goat serum. Human ASYN was detected using a specific rabbit anti-hASYN primary antibody (1:50000; Abcam, RRID:AB_2537217, MJFR1). After incubation with this antibody, sections were rinsed, kept for 1 hour in goat anti-rabbit biotinylated secondary antibody (1:200; Vector Laboratories, RRID:AB_231360) and, finally, treated with avidin-biotin–horseradish peroxidase complex (ABC Elite kit, Vector Laboratories). The brightfield signal was developed using a 3,3′-diaminobenzidine kit (Vector Laboratories).

Spreading of hASYN was assessed by counting the number of hASYN–immunoreactive axons in left (ipsilateral to the AAV injection side) sections from the pons (Bregma: −5.40 mm), midbrain (Bregma −4.60 mm) and forebrain (Bregma: −0.94 mm). All counts were performed by an investigator blinded to sample treatment using a Zeiss AXIO Observer microscope (Carl Zeiss) at high-magnification (63× Plan-Apochromat objective)^27^.

### Fluorescence microscopy and image analysis

Double immunofluorescence staining was performed on every 10th section of the medulla oblongata (MO)^27^. Samples were blocked in a solution containing 5% normal serum, 2% BSA and 0.5% Triton-X100 for 1 hour at room temperature. Tissue samples were incubated overnight in a solution containing both rabbit anti-hASYN (1:10000; Abcam, RRID:AB_2537217, MJFR1) and mouse anti-SynO2 (1:1000; Biolegend, RRID:AB_2632701). To detect immunofluorescence, sections were incubated for 1 hour in a solution containing donkey anti-rabbit and donkey anti-mouse antibodies conjugated with Alexa Fluor 555 and Alexa Fluor 647 (1:200; Invitrogen, RRID:AB_2762834 and RRID:AB_162542), respectively. Separate tissue sections were incubated first with rabbit anti-MJFR-14-6-4-2 (1:20000; Abcam, RRID:AB_2714215) or rabbit anti-SOD2 (1:1000; Cell Signaling, RRID:AB_2636921) and then with Alexa Fluor 555-conjugated donkey anti-rabbit secondary antibody (1:200; Abcam). These sections were also stained using fluorophore-conjugated rabbit anti-hASYN (1:400; Abcam, RRID:AB_2537217, MJFR1-Alexa 488).

After fluorescence staining and after collection of MO sections from mice injected with DHE, images (3 z-stacks, 280 nm step size) were acquired on a confocal microscope (Zeiss, LSM900) equipped with 63 x (N.A. 1.4) objectives, using ZEN 3.6 (Blue edition) software (Carl Zeiss, https://www.zeiss.com/microscopy/en/products/software/zeiss-zen.html RRID:SCR_013672). Intensities of hASYN, SynO2 and MJFR-14-6-4-2 immunostaining as well as intensity of the ox-DHE signal were measured using maximum intensity projection of z-stacks (ImageJ 1.54f). Background was subtracted using a rolling ball algorithm (radius of 50 pixels), and a mask was created with a constant threshold that allowed the specific detection of hASYN signal within neuronal cell bodies and neurites throughout the dorsal motor nucleus of the vagus (X^th^) nerve (DMnX). To perform measurements within perikarya, cell bodies were manually delineated. To isolate neurites, a second mask was created by excluding the delineated hASYN-positive cell bodies from the initial mask. Human ASYN, SynO2 and MJFR-14-6-4-2 intensities were measured separately within cell bodies, neurites or cell bodies plus neurites. DHE ethidium cation (ox-DHE) signal was measured only within perikarya^28^.

### In situ proximity ligation assay

Aggregated hASYN, nitrated hASYN and nitrated NDUFB8 [NADH dehydrogenase (ubiquinone) I beta subcomplex subunit 8] were detected using in situ proximity ligation assay (PLA), as previously described^27^. Aggregated hASYN PLA was performed using a “direct” detection method that involved conjugation of plus and minus PLA probes to anti-hASYN (Syn211; Millipore, RRID:AB_310817) using the Duolink Probemaker kit (Sigma-Aldrich). “Indirect” PLAs for nitrated hASYN and nitrated NDUFB8 were performed using probes already conjugation to secondary antibodies and provided in a PLA kit (Duolink, Sigma-Aldrich). Primary antibodies used for indirect PLA were: anti-3-nitrotyrosine (3-NT) (1:8000; Abcam, RRID:AB_942087) together with anti-hASYN (1:10000; Abcam, RRID:AB_2537217), or anti-3-NT (1:250) together with anti-NDUFB8 (1:300; Proteintech, RRID:AB_2150970). Following ligation and amplification steps, aggregated and nitrated hASYN signals were detected using a brightfield detection kit (Duolink, Sigma-Aldrich). Fluorescence detection of nitrated NDUFB8 was achieved using a Duolink Red Detection Kit (Sigma-Aldrich); sections were then stained with anti-hASYN (MJFR1) directly conjugated with Alexa 488 (1:500).

For quantification of hASYN/hASYN PLA, brightfield PLA dots were counted within DMnX neurons in MO sections at the level of the obex using the meander sampling option of Stereo Investigator (MBF Bioscience, RRID:SCR_018948, https://www.mbfbioscience.com/stereo-investigator). Brightfield hASYN/3-NT PLA dots were quantified stereologically in the DMnX of equally spaced MO sections (every tenth section between bregma −6.96 and −8.00 mm) using the optical dissector (Stereo Investigator, MBF Bioscience). Fluorescent NDUFB8/3-NT PLA signal was quantified withing hASYN-containing neurons in 3 MO sections between bregma −6.96 and −8.00 mm. The sections were imaged using a scanning confocal microscope (LSM 700), and maximum intensity projections were generated with the ZEN software (Zeiss). DMnX hASYN-immunoreactive neurons were delineated using Fiji (ImageJ version 2.1.0/1.53c, RRID:SCR_002285, http://fiji.sc) and, within these neurons, the area occupied by the PLA signal was quantified and expressed as the area fraction. Total intensity of the PLA signal was also quantified within the delineated hASYN-containing neurons. Cellular measurements of PLA area fraction and intensity were averaged for each animal.

### Statistical analysis

Statistical analyses were performed with Graph Pad Prism (version 10.0, RRID:SCR_002798, http://www.graphpad.com/). Normal distribution was checked with Shapiro-Wilk and Kolmogorov-Smirnov tests. Unpaired Student’s t-test or Mann-Whitney test was used for two-group comparisons. Two-way analysis of variance (ANOVA) was performed when different treatment groups were compared between control and transgenic mice. A specific post-hoc test (Tukey’s or Bonferroni’s test) was then used to identify significant differences. *P* values of less than 0.05 were considered statistically significant.

## Results

### ASYN aggregation and spreading in L444P/wt mice

The L444P mutation is one of the most common *GBA1* variants associated with increased PD risk and, for this reason, L444P-expressing mice represent a particularly valuable model for studying mutation-associated pathophysiological changes^3,29^. The experiments described here were performed in parallel using L444P/wt mice with heterozygous L444P expression and littermate controls. Measurements of GCase activity in whole brain homogenates showed a significant decrease of approximately 30% in samples from mutation-carrying as compared to control animals (Fig. 1a). Homogenates of the dMO were used for western blot analysis of ASYN content, which was found to be increased by approximately 30% in L444P/wt mice (Fig. 1b and c).

**Fig. 1.**
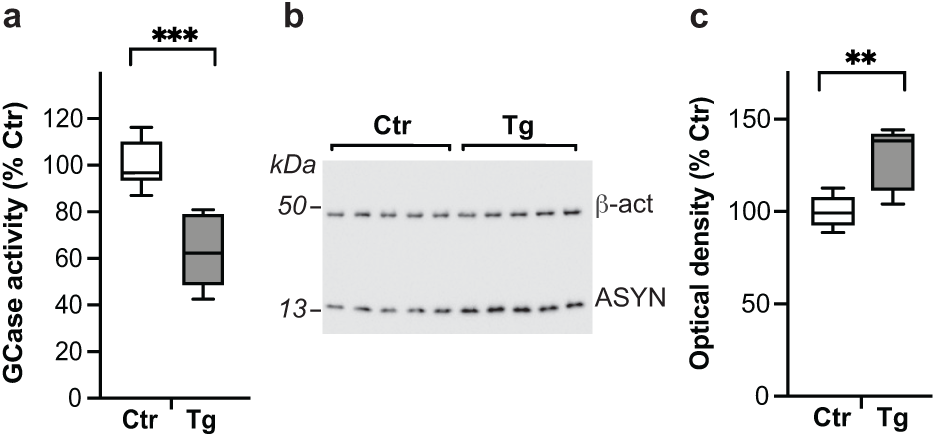
Features of heterozygous L444P transgenic mice. **a** GCase activity (expressed as percentage of the mean value in the control group) in brain homogenates from control (Ctr, *n* = 6) and L444P mutant (Tg, n = 7) mice. **b, c** Western blots from homogenates of the dMO showing immunoreactivity specific for ASYN and β-actin (**b**). Semiquantitative analysis of band optical density (calculated as ASYN/β-actin ratio and expressed as percentage of the mean value in the control group) in samples from control and L444P mutant mice (n = 5/group) (**c**). Plots show median, upper and lower quartiles, and maximum and minimum as whiskers. \*\**P* ≤0.01 and \*\*\**P* ≤ 0.001, Student’s t-test.

To investigate the effects of decreased GCase activity on overexpression-induced ASYN pathology, L444P/wt and control mice were treated with a single intravagal (left vagus nerve) injection of hASYN-delivering AAVs. In these animals, sections of the lower brainstem immunostained for hASYN and visualized with brightfield microscopy revealed robust transduction and expression of the exogenous protein in the dMO (Fig. 2a). In particular, consistent with earlier findings using this vagal paradigm, AAV-induced overexpression targeted neuronal cell bodies and neurites in the left DMnX as well as axonal projections that occupied both the left and right nucleus of the tractus solitarius^24–26^. The pattern and density of hASYN staining were similar in sections from control and L444P/wt mice (Fig. 2a). MO sections stained with anti-hASYN were also processed for fluorescent microscopy in order to visualize hASYN expression and quantify its intensity within DMnX neurons; data showed comparable levels of hASYN overexpression in samples from control and mutant animals (Fig. 2b and c). Western blot analyses of dMO homogenates were carried out at 2 weeks post-AAV injection, at a time when full expression of the exogenous protein is expected. Robust bands immunoreactive for total (mouse and human) ASYN were observed in samples from either L444P/wt or control mice; semiquantitative densitometric measurements revealed no significant differences in ASYN protein levels between the two groups of animals (Fig. 2d and e).

**Fig. 2.**
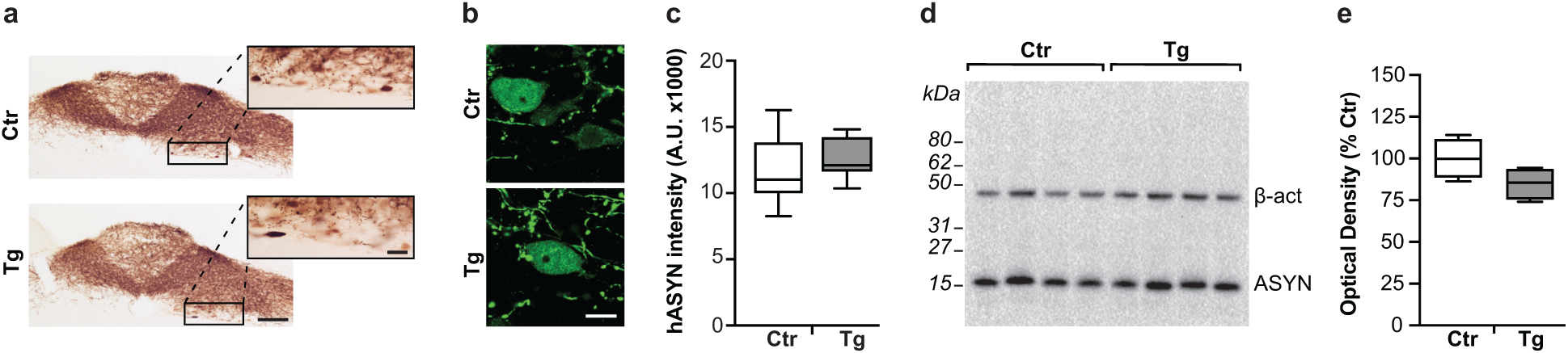
AAV-induced hASYN overexpression in the dMO. All mice received an intravagal injection of hASYN-delivering AAVs. **a** MO tissue sections were stained with a specific antibody against hASYN. Representative brightfield images of the dMO from a control (Ctr) and an L444P mutant (Tg) mouse. Boxes in the low-magnification images encompass areas of the DMnX that are also shown at higher magnification. Scale bars: 250 μm (low magnification) and 20 μm (higher magnification). **b, c** Representative fluorescent images of neurons in the left (ipsilateral to the AAV injection) DMnX stained with anti-hASYN. Scale bar: 10 μm (**b**). Fluorescent intensity (expressed as arbitrary units, A.U.) of hASYN-positive neurons in the left DMnX of control (n = 6) and L444P mutant (n = 8) mice (**c**). **d, e** Western blots from homogenates of the dMO showing immunoreactivity specific for total (rodent plus human) ASYN and β-actin (**d**). Semiquantitative analysis of band optical density (calculated as ASYN/β-actin ratio and expressed as percentage of the mean value in the control group) in samples from control and L444P mutant mice (n = 4/group) (**e**). Plots show median, upper and lower quartiles, and maximum and minimum as whiskers.

Overexpression-induced ASYN aggregation was assessed in medullary tissue sections collected from control and L444P/wt mice at 6 weeks post-AAV treatment and stained with combinations of 3 different antibodies. One of these antibodies (MJFR1) recognizes both monomeric and aggregated hASYN, whereas the other two reagents (SynO2 and MJFR-14-6-4-2) display greater avidity for aggregated forms of the protein^30–32^. A first set of sections was double-stained with MJFR1 and SynO2 and processed for semiquantitative intensity measurements in the whole DMnX or within DMnX cell bodies and neurites. Both microscopy observations and quantitative analyses showed that immunoreactivity for MJFR1 was comparable between control and L444P/wt samples, whereas SynO2 intensity was significantly increased in sections from mutant animals (Fig. 3a and b and Supplementary Fig. 2). Similar findings were obtained when a second set of samples was double-labeled with MJFR1 and MJFR-14-6-4-2; MJFR1 immunoreactivity remained unchanged while MJFR-14-6-4-2 labeling was higher in the DMnX of L444P/wt as compared to control mice (Supplementary Fig. 3). Taken together, these data suggest that, while levels of total (monomeric plus aggregated) hASYN are not affected by L444P expression, the proportion of aggregated protein is enhanced in the presence of mutated GCase. L444P-induced ASYN aggregation was then further assessed using a specific, highly sensitive PLA, detecting self-interactions between hASYN molecules (hASYN/hASYN PLA) ^25,33^. Following PLA signal amplification, chromogenic dots were observed in the dMO of all AAV-injected mice (Fig. 3c). The number of these dots was much greater, however, in tissue from mutant animals, in which dot counts revealed a 5-fold increase within DMnX cell bodies (Fig. 3c and d).

**Fig. 3.**
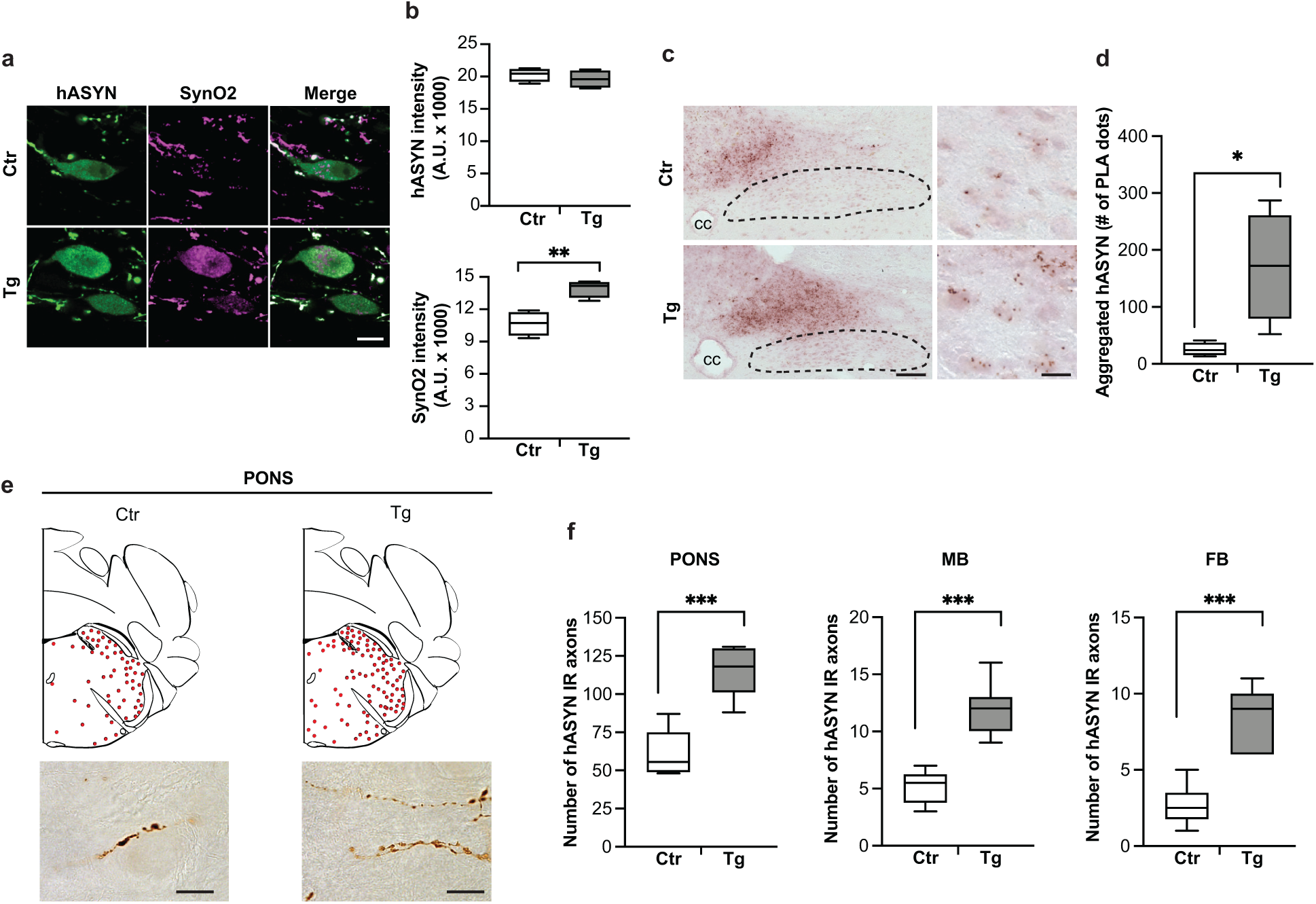
L444P-induced aggregation and spreading of hASYN. All mice received an intravagal injection of hASYN-delivering AAVs. **a, b** MO tissue sections were double-stained with anti-hASYN and an antibody, SynO2, that reacts with aggregated ASYN species. Representative fluorescent images of neurons in the left (ipsilateral to the AAV injection) DMnX of a control (Ctr) and an L444P mutant (Tg) mouse. Scale bar: 10 μm (**a**). Fluorescent intensity (expressed as arbitrary units, A.U.) of hASYN immunoreactivity was measured in the left DMnX; fluorescent SynO2 intensity was measured within hASYN-containing DMnX neurons (both cell bodies and neurites); samples were obtained from control and L444P mutant mice (n = 4/group) (B). **c, d** MO sections from AAV-injected mice were processed for PLA using probes that specifically detect aggregated hASYN. Representative low magnification images of the left-middle dMO in which the central canal (cc) is indicated and the DMnX is delineated with dashed lines. Higher magnification images show neurons in the left DMnX. Scale bars: 100 μm (low magnification) and 10 μm (higher magnification) (**c**). The number of neuronal PLA dots was counted in the left DMnX of control and L444P mutant mice (n = 4/group) mice (**d**). **e, f** Tissue sections were stained with anti-hASYN. Schematic plots of the distribution of hASYN-labelled axons (each red dot represents one of these axons) in the left pons. Representative images below show hASYN-immunoreactive axons in the left pons. Scale bars: 40 μm (**e**). The number of hASYN-positive axons was counted in tissue sections of the left pons, midbrain (MB) and forebrain (FB) from control and L444P mutant mice (n ≥ 6/group) (**f**). Plots show median, upper and lower quartiles, and maximum and minimum as whiskers. \**P* ≤ 0.05, \*\**P* ≤ 0.01 and \*\*\**P* ≤ 0.001, Student’s t-test (**b** and **f**) and Mann-Whitney U test (**d**).

When overexpressed within “donor” neurons in the DMnX, hASYN can be transferred into “recipient” axons projecting into the dMO and, through this route, spread toward higher brain regions^24–26^. This spreading process was compared in control *vs.* L444P/wt mice sacrificed at 6 weeks after vagal AAV administration. Tissue sections of the pons, midbrain and forebrain were immunostained with anti-hASYN and, in each section, the number of positive axons containing the exogenous protein was counted. Results showed that, while dystrophic axons loaded with hASYN were observed in pontine tissue from either control or L444P/wt animals, their counts were significantly increased in samples from mutant mice (Fig. 3e and f). Similarly, a higher number of hASYN-positive fibers was counted in sections from the midbrain and forebrain of L444P/wt mice, further supporting the conclusion that heterozygous L444P expression was associated with enhanced interneuronal transfer and more pronounced caudo-rostral brain spreading of hASYN (Fig. 3f).

### Vulnerability to Ox/Nt stress conferred by the L444P mutation

GCase mutations have been shown to induce Ox stress, and Ox stress may exacerbate ASYN pathology^28,34^. Based on these considerations, the next set of experiments was aimed at determining whether enhanced vulnerability to Ox stress was a feature of L444P-expressing neurons. Formation and accumulation of ROS was compared in hASYN-overexpressing DMnX neurons from L444P/wt *vs.* control mice. Following vagal AAV-treatment, mice were kept for 6 weeks and then injected with the superoxide indicator DHE shortly before the time of sacrifice. Upon reacting with superoxide, DHE is converted to the fluorescent ox-DHE that can be visualized and quantified using fluorescence microscopy. Ox-DHE labelling was detected within hASYN-containing DMnX neurons in MO sections from both control and L444P/wt mice (Fig. 4a). Mutation-expressing neurons from the latter group were characterized, however, by a more widespread and significantly more intense fluorescent signal (Fig. 4a and b).

**Fig. 4.**
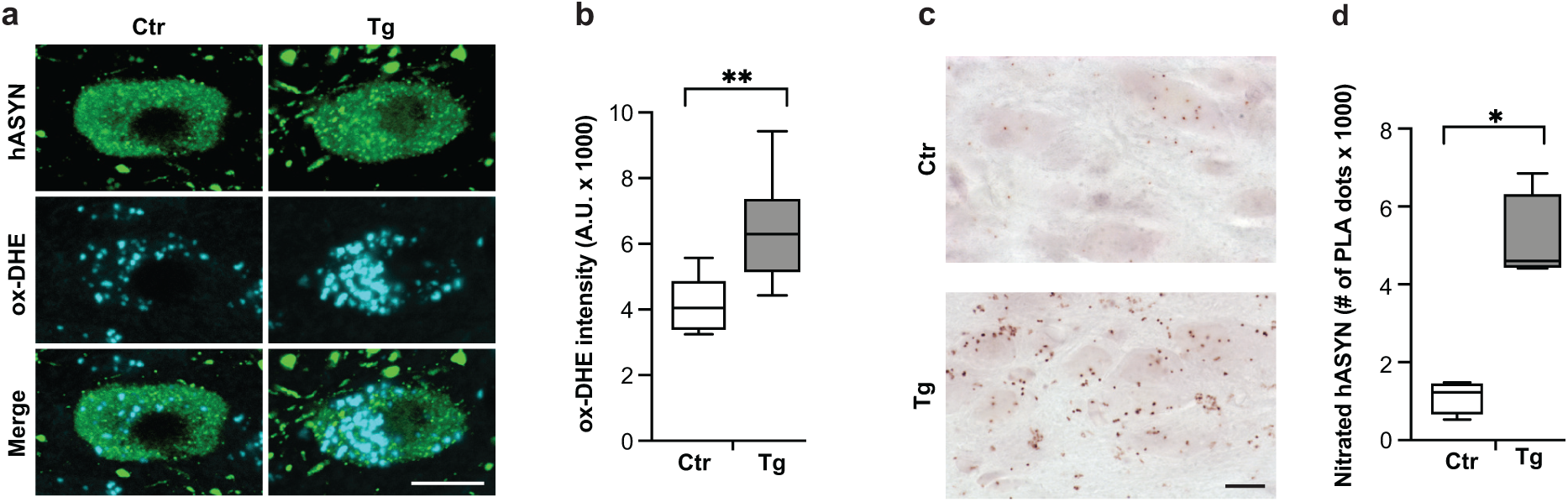
L444P-induced Ox and Nt stress. All mice received an intravagal injection of hASYN-delivering AAVs. **a, b** At the time of sacrifice, mice were injected subcutaneously with DHE. Representative images of fluorescent ox-DHE, a superoxide indicator, within hASYN-containing neurons in the left (ipsilateral to the AAV injection) DMnX of a control (Ctr) and an L444P mutant (Tg) mouse. Scale bar: 10 μm (**a**). Fluorescent ox-DHE intensity (expressed as arbitrary units, A.U.) within hASYN-containing neurons in the left DMnX of control (n = 6) and L444P mutant (n = 8) mice (**b**). **c, d** MO sections were processed for PLA using probes that specifically detect nitrated hASYN. Representative images of the left DMnX. Scale bars: 10 μm (**c**). The number of neuronal PLA dots was counted in the left DMnX of control and L444P mutant mice (n = 4/group) (**d**). Plots show median, upper and lower quartiles, and maximum and minimum as whiskers. \**P* ≤ 0.05 and \*\**P* ≤ 0.01, Student’s t-test (**b**) and Mann-Whitney U test (**d**).

Increased ROS production can be paralleled by the accumulation of reactive nitrogen species (RNS), resulting in Nt modifications of cellular macromolecules and, ultimately, Nt stress. Proteins containing tyrosine residues, including ASYN, can be modified by Nt reactions, and detection of nitrated proteins is not only a marker of Nt stress but may also indicate structural and behavioural changes of the modified proteins. To test occurrence and severity of Nt stress in the absence or presence of mutated GCase, accumulation of nitrated hASYN was assessed in MO sections from AAV-injected control and L444P/wt mice. Tissue samples were processed using a PLA specifically designed to detect nitrated hASYN^28^. MO sections were incubated first with a pair of primary antibodies, namely anti-hASYN and anti-3-NT, and then with secondary antibodies conjugated with PLA probes (hASYN/3-NT PLA). Following PLA signal amplification, bright-field microscopy showed specific chromogenic dots in the dMO of all hASYN-overexpressing animals; the density and intensity of the dots were apparently enhanced, however, in sections from L444P/wt mice (Fig. 4c). Quantification of this nitrated hASYN signal was carried out by PLA dot counts, and comparative analyses revealed a marked, 5-fold signal increase within DMnX neurons from mutation-carrying animals (Fig. 4d). Taken together, these findings indicate that neuronal L444P expression promotes ROS production and ROS/RNS reactions, thus rendering DMnX neurons highly vulnerable to Ox and Nt stress and ensuing pathology.

### Relationship between Ox stress and L444P-induced ASYN pathology

Results showing exacerbated ASYN pathology and greater susceptibility to Ox/Nt stress in L444P/wt mice raised the question of whether a causal relationship links mutation-induced ROS/RNO production to ASYN aggregation and spreading. To address this question, rescue experiments were designed in which neuronal ROS burden was counteracted by overexpression of SOD2, an antioxidant enzyme particularly important for the scavenging of mitochondria-derived superoxide. AAVs delivering SOD2 DNA or empty vectors lacking protein coding sequence were co-injected with hASYN-AAVs into the left vagus nerve of animals divided into the following 4 experimental groups: control mice co-injected with (1) hASYN- and empty-AAVs, or (2) hASYN- and SOD2-AAVs, and L444P/wt mice treated with (3) hASYN- and empty-AAVs, or (4) hASYN- and SOD2-AAVs. Animals were sacrificed at 6 weeks after AAV injections. To verify correct co-transduction and overexpression of both hASYN and SOD2, MO sections from mice belonging to the 4 different experimental groups were double-stained with anti-hASYN and anti-SOD2 and visualized using fluorescent microscopy. Immunofluorescence specific for hASYN characterized the dMO of all injected mice, whereas a marked increase in SOD2 reactivity was only evident within hASYN-containing DMnX neurons from control and mutant animals that had received combined injections of hASYN/SOD2-AAVs (Fig. 5a). Homogenates of the dMO were processed for western blot analysis to verify that total (endogenous and human) ASYN levels were comparable in the different experimental groups; indeed, semiquantitative measurements of band optical density showed no differences between control and L444P/wt mice and between animals injected with either hASYN/empty- or hASYN/SOD2-AAVs (Fig. 5b and c).

**Fig. 5.**
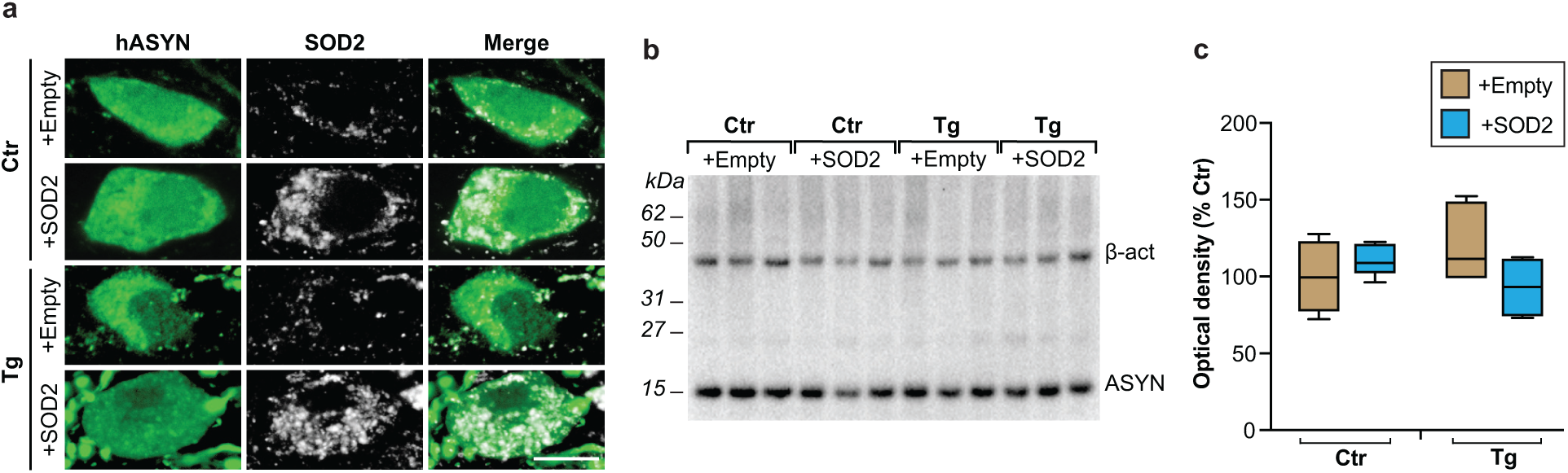
Co-overexpression of hASYN and SOD2. Control (Ctr) and L444P mutant (Tg) mice were all injected intravagally with hASYN-delivering AAVs. Groups of animals also received, together with hASYN-AAVs, empty vectors lacking protein coding (+empty) or AAVs delivering SOD2 DNA (+SOD2). **a** MO tissue sections were double-stained with anti-hASYN and anti-SOD2. Representative fluorescent images of neurons in the left (ipsilateral to the AAV injections) DMnX. Scale bar: 10 μm. **b, c** Western blots from homogenates of the dMO showing immunoreactivity specific for total (rodent plus human) ASYN and β-actin (**b**). Semiquantitative analysis of band optical density (calculated as ASYN/β-actin ratio and expressed as percentage of the mean value in the control group injected with hASYN/empty AAVs) in samples from control and L444P mutant mice co-injected with either hASYN/empty(light brown)- or hASYN/SOD2(azure blue)-AAVs (n ≥ 4/group) (**c**). The plot shows median, upper and lower quartiles, and maximum and minimum as whiskers.

In the next set of experiments, samples from the 4 groups of control and transgenic mice with or without SOD2 overexpression were used to asses ASYN pathology. To detect and compare aggregate pathology, MO sections were double-stained with anti-hASYN and anti-SynO2. Examination of fluorescent images with confocal microscopy showed comparable immunoreactivity for hASYN in samples from all animals regardless of their treatment (Fig. 6a). Intraneuronal SynO2 labelling, which more specifically detect aggregated ASYN species, was overtly enhanced in the left DMnX of transgenic mice injected with hASYN/empty-AAVs; it was instead relatively weak and unchanged in the DMnX of all other animals, including L444P/wt mice treated with hASYN/SOD2-AAVs (Fig. 6a). Quantitative assessment of labelling intensities in the whole DMnX as well as within DMnX cell bodies or neurites confirmed the results of the microscopy observations, showing a significantly increased immunoreactivity only for SynO2 and only in the left DMnX of hASYN/empty-AAV-injected transgenic mice (Fig. 6b and Supplementary Fig. 4). Protein aggregation was also evaluated using hASYN/hASYN PLA. An increased number of neuronal chromogenic PLA dots characterized the left DMnX in MO sections from L444P/wt mice treated with hASYN/empty-AAVs, whereas the PLA signal in transgenic mice overexpressing SOD2 remained low and comparable to the signal seen in control animals (Fig. 6c and d). Thus, overexpression of SOD2 effectively prevented the accumulation of aggregated ASYN that was otherwise induced by the mutated GCase.

**Fig. 6.**
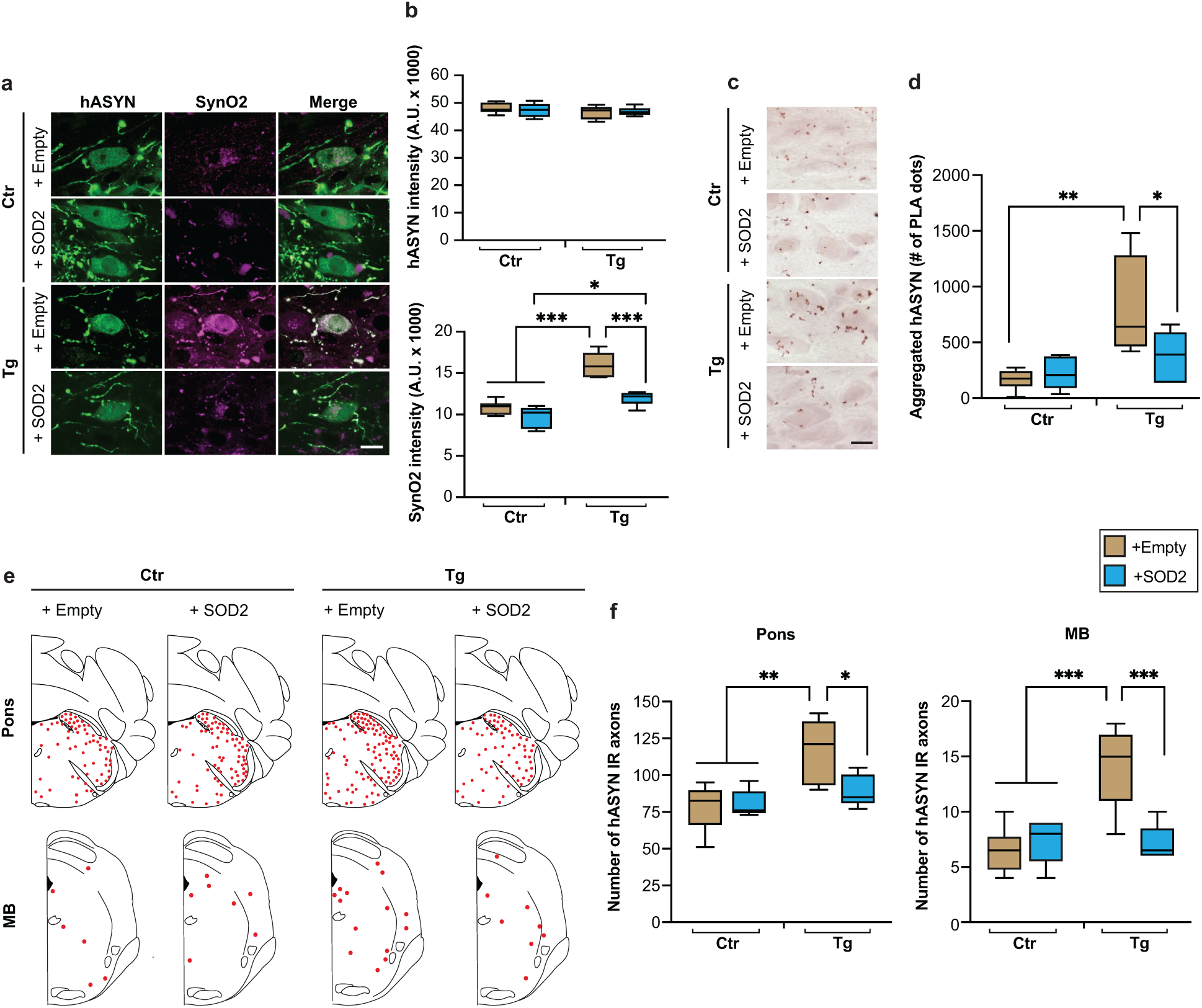
Effects of SOD2 transduction on L444P-induced hASYN aggregation and spreading. Control (Ctr) and L444P mutant (Tg) mice were all injected intravagally with hASYN-delivering AAVs. Groups of animals also received, together with hASYN-AAVs, empty vectors lacking protein coding (+empty) or AAVs delivering SOD2 DNA (+SOD2). **a, b** MO tissue sections were double-stained with anti-hASYN and anti-SynO2. Representative fluorescent images of neurons in the left (ipsilateral to the AAV injections) DMnX. Scale bar: 10 μm (**a**). Fluorescent intensity (expressed as arbitrary units, A.U.) of hASYN immunoreactivity was measured in the left DMnX; fluorescent SynO2 intensity was measured within hASYN-containing DMnX neurons (both cell bodies and neurites); samples were obtained from control and L444P mutant mice co-injected with either hASYN/empty(light brown)- or hASYN/SOD2(azure blue)-AAVs (n ≥ 5/group) (**b**). **c, d** MO sections were processed for PLA using probes that specifically detect aggregated hASYN. Representative images of the left DMnX. Scale bar: 10 μm (**c**). The number of neuronal PLA dots was counted in the left DMnX of control and L444P mutant mice co-injected with either hASYN/empty- or hASYN/SOD2-AAVs (n ≥ 4/group) (**d**). **e, f** Tissue sections were stained with anti-hASYN. Schematic plots of the distribution of hASYN-labelled axons (each red dot represents one of these axons) in the left pons and midbrain (M**B**) (**e**). The number of hASYN-positive axons was counted in tissue sections of the left pons and MB from control and L444P mutant mice co-injected with either hASYN/empty- or hASYN/SOD2-AAVs (n ≥ 5/group) (**f**). Plots show median, upper and lower quartiles, and maximum and minimum as whiskers. \**P* ≤ 0.05, \*\**P* ≤ 0.01 and \*\*\**P* ≤ 0.001, Tukey’s (**b** and **f**) and Bonferroni (**d**) tests.

To determine if SOD2 transduction also affected L444P-induced hASYN spreading (see Fig. 3e and f), sections of the pons and midbrain were obtained from control and transgenic animals that received intravagal injections of hASYN/empty- or hASYN/SOD2-AAVs. After staining with anti-hASYN, hASYN-positive axons were visualized and quantified. Representative schematic plots of the number and distribution of immunoreactive axons in pontine and midbrain sections show no overt distribution differences among samples from the 4 treatment groups (Fig. 6e). Counts of hASYN-containing axons revealed significant changes, but only in L444P/wt mice treated with hASYN/empty AAVs; in this group of animals, the number of hASYN-positive axons was augmented by approximately 50% and 100% in the pons and midbrain, respectively (Fig. 6e and f). Lack of increased spreading in transgenic mice injected with hASYN/SOD2 AAVs indicates that enhanced ROS scavenging within donor DMnX neurons is enough to reverse the effects of mutated GCase. These data are also consistent with the interpretation that Ox stress plays a critical mechanistic role in L444P-induced interneuronal hASYN transfer.

### Nt damage within L444P-expressing neurons

Evidence of accumulation of nitrated hASYN in the DMnX of L444P/wt mice (see Fig. 4c and d) prompted experiments that assessed whether this buildup was affected by SOD2 overexpression. Tissue sections of the MO were obtained from control and L444P/wt mice treated with hASYN/empty- or hASYN/SOD2-AAVs; they were then processed for hASYN/3-NT PLA. A significantly increased PLA signal was observed and quantified in the left DMnX of transgenic as compared to control mice injected with hASYN/empty-AAVs (Fig. 7a and b). This L444P-induced nitrated hASYN burden was completely reversed, however, when overexpression of hASYN was paralleled by SOD2 transduction in mice injected with hASYN/SOD2-AAVs (Fig. 7a and b). Thus, superoxide dismutation not only counteracted ROS buildup and injury but also effectively prevented ROS/RNS reactions leading to ASYN nitration.

**Fig. 7.**
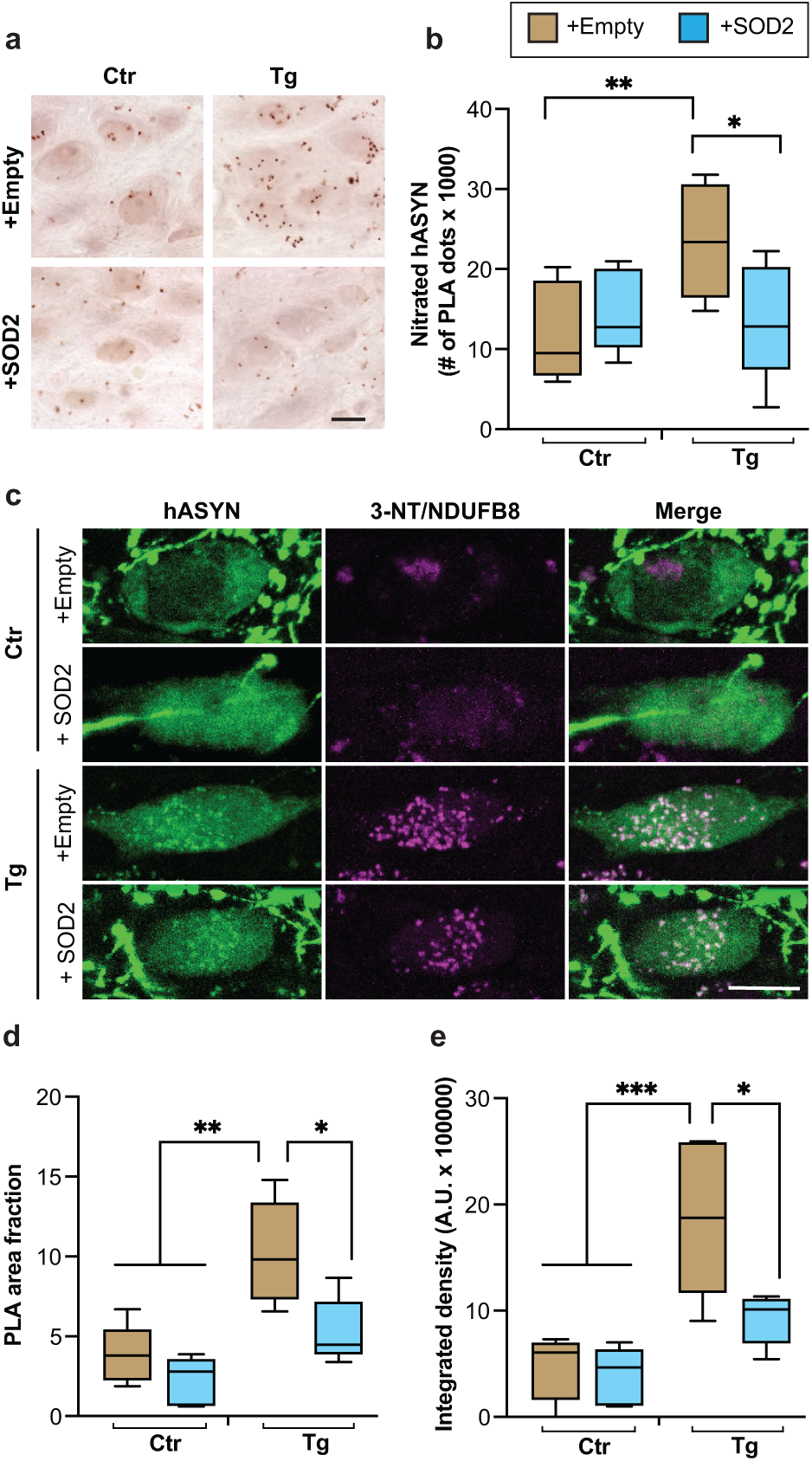
L444P-induced nitrative protein modifications. Control (Ctr) and L444P mutant (Tg) mice were all injected intravagally with hASYN-delivering AAVs. Groups of animals also received, together with hASYN-AAVs, empty vectors lacking protein coding (+empty) or AAVs delivering SOD2 DNA (+SOD2). **a, b** MO tissue sections were processed for detection of nitrated hASYN using PLA. Representative images of PLA chromogenic dots in the left (ipsilateral to the AAV injections) DMnX. Scale bar: 10 μm (**a**). The number of neuronal PLA dots was counted in the left DMnX in samples from Ctr and Tg mice co-injected with either hASYN/empty(light brown)- or hASYN/SOD2(azure blue)-AAVs (n ≥ 5/group) (**b**). **c-e** MO sections were processed for PLA detecting nitrated NDUFB8 and also labelled with anti-hASYN. Representative fluorescent images of neurons in the left (ipsilateral to the AAV injections) DMnX. Scale bar: 10 μm (**c**). PLA area fraction (**d**) and integrated density of the PLA signal (**e**) were quantified in fluorescent samples from Ctr and Tg mice co-injected with either hASYN/empty- or hASYN/SOD2-AAVs (n = 5/group). Plots show median, upper and lower quartiles, and maximum and minimum as whiskers. \**P* ≤ 0.05, \*\**P* ≤ 0.01 and \*\*\**P* ≤ 0.001, Bonferroni (**b**) and Tukey’s (**d** and **e**) tests.

The occurrence of ROS/RNS reactions would be expected to modify other proteins besides ASYN. Furthermore, the ability of mitochondrial SOD2 to reverse L444P-induced pathological changes suggests an important toxic role of mitochondrial ROS production and raises the possibility that mitochondrial proteins may be highly susceptible to Ox/Nt modifications. To test these possibilities, Nt modifications of mitochondrial complex I subunit NDUFB8 were examined and compared in control and L444P/wt mice that overexpressed hASYN in the absence or presence of concurrent SOD2 transduction. NDUFB8 nitration was detected and quantified using a specific PLA that involved incubations of MO sections first with anti-NDUFB8 and anti–3-NT and then with oligonucleotide-labelled secondary antibodies and hybridizing connector oligonucleotides. Tissue sections were also stained with anti–hASYN and analysed by confocal microscopy. Following treatment with hASYN/empty-AAVs, detection of the PLA signal showed a marked difference in specimens from L444P/wt as compared to control mice, with the former displaying enhanced nitrated NDUFB8 fluorescence within hASYN-containing DMnX neurons (Fig. 7c). This effect was not observed when animals were injected with hASYN/SOD2-AAVs since, under this experimental condition, a relatively weak and comparable PLA signal characterized hASYN-expressing neurons in the DMnX of either control or transgenic mice (Fig. 7c). These microscopy observations were confirmed by measurements of the intracellular area occupied by PLA dots and quantification of PLA signal intensity; with both assays, a significant increase was found only in samples from L444P/wt mice injected with hASYN/empty-AAVs (Fig. 7d and e). Enhanced NDUFB8 nitration in the DMnX of L444P mice not only provides further evidence of mutation-induced Nt stress but also supports a mechanism of toxicity that, by damaging mitochondrial proteins, may contribute to neuronal dysfunction and pathology. Another important finding of these experiments was the reversal of L444P-induced NDUFB8 nitration in neurons overexpressing SOD2 (Fig. 7d and e). Thus, strategies aimed at preventing superoxide accumulation may be sufficient also to alleviate ROS/RNS reactions and ultimately to protect against the deleterious consequences of mitochondrial protein nitration.

## Discussion

Results of this study address several important questions concerning the relationship between *GBA1* mutations and vulnerability to ASYN pathology. First, experiments and analyses were designed to compare ASYN aggregation and spreading between mice carrying a heterozygous L444P mutation of GCase *vs.* littermate controls. Pathological changes specifically targeted the dMO and were induced by overexpression of hASYN after treatment with an intravagal injection of hASYN-delivering AAVs^24,25^. Data unequivocally revealed significantly more severe ASYN pathology in AAV-injected L444P/wt mice. In these transgenic animals, increased ASYN aggregation was detected in DMnX-containing MO sections stained with antibodies that react with self-assembled forms of the protein^30,31^. Enhanced aggregate pathology was also evident in tissue samples processed for PLA using probes highly specific for non-monomeric hASYN^25,33^. A separate set of analyses revealed a mutation-induced exacerbation of ASYN brain spreading that proceeded caudo-rostrally from the dMO and was quantified in pontine, midbrain and forebrain regions. Evidence of more severe ASYN pathological burden in the DMnX of L444P/wt mice bears significant translational implications. The DMnX is a primary site affected by ASYN aggregate pathology at early stages of PD development^35^. It may also play a key role in the spreading of ASYN pathology that, in parkinsonian brains, often advances following a stereotypical pattern from the lower brainstem toward higher brain regions^36^. In PD, ASYN inclusions are not only features of neuronal brain cells but have also been detected within neurons of the peripheral nervous system (PNS) ^37,38^. Long-distance transfer of pathogenic ASYN lesions from PNS to brain neurons and *vice versa* may contribute to pathology progression and would likely occur *via* vagal projections that, from the DMnX, reach peripheral tissues throughout the body^26,36,39^. Based on these considerations, our present findings reveal that, in individuals carrying *GBA1* mutations (in particular, the L444P mutation), enhanced vulnerability of DMnX neurons to ASYN aggregation and spreading may play an important role in early development and subsequent advancement of PD pathology; it could also facilitate peripheral-to-central or central-to-peripheral transmission of the pathological process.

In the dMO of our naïve, untreated L444P mutant mice, reduced GCase activity was found to be associated with increased ASYN levels, consistent with the results of earlier *in vitro* and *in vivo* studies showing an inverse relationship between enzyme activity and ASYN burden^11,17,40,41^. Interestingly, however, when ASYN levels were assessed both immunohistochemically and biochemically after AAV injections, no significant changes were observed in the dMO of mutant as compared to control animals. Specific mechanisms underlying these different findings in the absence or presence of AAV administration remain unclear, although it is quite possible that the robust hASYN overexpression after AAV treatment may overshadow the increase in neuronal ASYN content otherwise associated with the mutated GCase. The observation of similar ASYN levels in overexpressing control and mutant mice raises another important consideration. It suggests that exacerbation of ASYN aggregation and spreading in AAV-injected transgenic animals was unlikely to be a mere consequence of differences in neuronal ASYN content. Other potential pathogenetic mechanisms were therefore hypothesized and investigated in the subsequent part of this study.

Several lines of earlier experimental and pathological evidence prompted our interest in the relationship between ASYN pathology and Ox stress in the DMnX of L444P/wt mice. First, cholinergic DMnX neurons have been shown to be highly susceptible to Ox stress, especially in the presence of enhanced ASYN expression^28^. Furthermore, Ox stress within DMnX neurons is capable of exacerbating ASYN aggregation and caudo-rostral ASYN spreading^27,28^. Loss of GCase activity, either chemically induced or associated with gene mutations, can cause Ox stress in a variety of *in vitro* systems, such as SHSY-5Y cells treated with conduritol-β-epoxide or primary hippocampal neurons isolated from L444P/wt mice^19,34^. A relationship between *GBA1* mutations and Ox stress is further supported by observations in humans. In particular, increased Ox stress has been reported within pyramidal neurons in the anterior cingulate cortex of post-mortem brain specimens from parkinsonian patients carrying *GBA1* mutations^19^. Results of the present study revealed increased neuronal accumulation of both ROS and RNS within hASYN-overexpressing DMnX neurons of L444P/wt mice. Since these neurons were also more vulnerable to ASYN pathology, these data provided first evidence of an association between DMnX expression of mutant GCase, Ox/Nt stress and ASYN aggregation and spreading. Ox stress has long been hypothesized to be a key mechanism underlying the susceptibility of discrete neuronal populations, such as dopaminergic cells in substantia nigra pars compacta and cholinergic neurons in the DMnX, to PD pathogenetic processes^28,42^. The present data suggest therefore that, at least in the DMnX, this “intrinsic” vulnerability to Ox stress-induced pathology may be significantly augmented in neurons expressing a GCase mutation.

Evidence of a parallel increase in ROS/RNS production and ASYN aggregation and spreading in L444P/wt mice suggests, but does not necessarily prove, a causal relationship between Ox stress and L444P-induced ASYN pathology. A separate set of experiments was therefore designed to counteract ROS accumulation and consequent Ox stress and to determine whether this rescue strategy successfully alleviated ASYN aggregation and spreading. Results of earlier investigations suggest that impaired or damaged mitochondria may represent primary sources of enhanced ROS production in cells with lower GCase activity as well as brain tissue samples from PD patients with *GBA1* mutations^19,34^. For this reason, the rescue approach used for these experiments was to enhance neuronal expression of the mitochondrial superoxide scavenging enzyme SOD2. SOD2- or empty-AAVs were injected together with hASYN-AAVs into the vagus nerve of control and L444P/wt mice. Immunohistochemical and biochemical analyses verified that levels of ASYN protein were unchanged between all groups of animals, regardless of their treatment with SOD2/hASYN-AAVs or empty/hASYN-AAVs. Further analyses verified that efficient co-transduction and co-expression of SOD2 and hASYN occurred only in the DMnX of control and transgenic mice treated with SOD2/hASYN-AAVs.

ASYN pathology was then assessed and compared among these different treatment conditions. Results showed that, in the absence of SOD2 overexpression (i.e., after treatment with empty/hASYN-AAVs), Ox stress and ASYN pathology were significantly exacerbated in L444P/wt mice. Quite remarkably, however, these effects of mutated GCase were completely reversed by SOD2 overexpression; indeed, following treatment with SOD2/hASYN-AAVs, levels of nitrated hASYN as well as burden of hASYN aggregation remained comparable in the DMnX of control and mutant mice. Furthermore, in the presence of SOD2 transduction, caudo-rostral hASYN spreading was no longer exacerbated by L444P expression, and the number of hASYN-positive axons was significantly reduced in pontine and midbrain sections from transgenic animals. Taken together, these data demonstrated that ROS scavenging effectively protected against L444P-induced ASYN pathology, providing compelling evidence for a mechanistic role of Ox stress in enhancing neuronal vulnerability to aggregation and spreading in the presence of a GCase mutation. Another interesting observation arising from the results of these experiments was that SOD2 overexpression, while reversing the enhanced pathology associated with L444P expression, did not completely abolish ASYN aggregation and spreading in transgenic mice, nor did it significantly affect protein assembly and transfer in control animals. It is conceivable that the residual pathology in L444P/wt mice as well as ASYN aggregation and spreading in control animals may be triggered by mechanisms other than Ox stress and therefore unresponsive to ROS scavenging. Alternatively, or in addition, ASYN pathology in mice treated with SOD2/hASYN-AAVs may still be a consequence of Ox stress; this stress, however, may involve production and accumulation of ROS from sources other than mitochondria (e.g., the endoplasmic reticulum) that would be unaffected by SOD2 overexpression.

Under Ox stress conditions, reaction of superoxide with nitric oxide generates highly unstable peroxynitrite anions that can in turn react with tyrosine residues of proteins and modify their structure and properties^43^. The pathophysiological relevance of these reactions is underscored by two main considerations: ASYN, with its 4 tyrosine residues, can be targeted by Nt modifications and, indeed, extensive and widespread ASYN nitration is a feature of Lewy inclusions in human synucleinopathies^44,45^. Data shown in this study are consistent with the occurrence of toxic reactions involving ROS and RNS that are triggered by hASYN overexpression and markedly enhanced by the L444P GCase mutation. Intraneuronal accumulation of nitrated ASYN was a clear marker of these reactions and strictly paralleled the severity of Ox stress and ASYN pathology. More pronounced ROS formation and more severe ASYN aggregation and spreading were associated with enhanced ASYN nitration in the brain of L444P mice. On the other hand, when treatment of these transgenic animals with SOD2-AAVs counteracted the superoxide burden and mitigated ASYN pathology, levels of nitrated ASYN were also significantly reduced. Results of earlier investigations provide further clues relevant to the interpretation of our present findings. Protein nitration has been shown to modulate ASYN’s ability to self-assembly^46,47^. Furthermore, once nitrated, ASYN appears to acquire greater mobility that promotes its intercellular exchanges^28^. Thus, accumulation of nitrated ASYN, as observed in the DMnX of transgenic L444P mice, may not only be a marker of pathological processes but could itself contribute to more prominent aggregate pathology and more advanced neuron-to-neuron brain spreading.

A final set of experiments of this study aimed at further exploring the effects of mutated GCase on protein nitration and, in particular, the nitration of mitochondrial protein. Results of experiments in SOD2-overexpressing mice pointed to mitochondria as important sources of L444P-induced superoxide formation. They also raised the possibility that, being close to sites of ROS production, mitochondrial proteins would be highly susceptible to Nt modifications. To test this hypothesis, nitration of a key mitochondrial protein, namely complex I subunit NDUFB8, was assessed using a specific PLA and compared in control and mutant mice that overexpressed hASYN in the absence or presence of SOD2 transduction. Neuronal levels of nitrated NDUFB8 were found to be significantly augmented in the DMnX of L444P/wt animals injected with empty/hASYN-AAVs; they instead remained unchanged between control and transgenic mice when both hASYN and SOD2 were overexpressed after treatment with SOD2/hASYN-AAVs. Thus, in neurons with mutated GCase, increased mitochondrial ROS burden and consequent reactions between ROS and RNS lead to an enhanced nitration of mitochondrial proteins. At least two additional considerations underscore the toxic/pathological relevance of L444P-induced NDUFB8 nitration. First, the NDUFB8 subunit plays a critical role in mitochondrial complex I assembly, stability and function that could be severely impaired by Nt reactions^48,49^. Second, an intriguing relationship between mitochondrial complex I and GCase has recently been reported. GCase protein was found to be imported from the cytosol into mitochondria where it modulated complex I integrity and function through mechanisms that were altered by PD-linked GCase mutations, such as L444P-GCase^50^. Taken together, current findings and results of this earlier investigation suggest a toxic, possibly self-amplifying mechanism that could contribute to pathological changes in the presence of mutated GCase. Mutation-induced complex I impairment may enhance ROS generation, stimulate ROS/RNS reactions and cause nitration of mitochondrial proteins, including NDUFB8; this protein nitration could in turn worsen complex I dysfunction, aggravate Ox and Nt stress and promote ASYN aggregation and spreading.

In summary, experimental evidence reported here provides new mechanistic clues into the relationship between a common GCase mutation, neuronal vulnerability to PD pathogenetic processes and development of ASYN lesions. In a brain region, namely the DMnX, highly susceptible to ASYN pathology and involved in propagation of this pathology, neuronal expression of mutated GCase was associated with enhanced ASYN assembly and spreading. Data proved a mechanistic link between expression of mutated GCase, mitochondrial ROS burden, Ox/Nt stress and exacerbation of aggregation and interneuronal brain transfer of ASYN. Rescue experiments demonstrated the feasibility of a protective intervention that, in the presence of a GCase mutation, may avert pathology conversion by counteracting ROS and RNS burden and/or preventing the accumulation of toxic products of Ox/Nt reactions. These products, as suggested by results of this study, may include nitrated ASYN and oxidatively/nitratively modified mitochondrial proteins.

## Acknowledgements

This work was supported by grants from the EU Joint Programme**-**Neurodegenerative Disease Research (JPND 01ED2005B, GBA-PaCTS) (DDM and AHVS). DDM, AU, PLV, ES and AHVS were in part funded by Aligning Science Across Parkinson’s (ASAP-000420) through the Michael J. Fox Foundation for Parkinson’s Research (MJFF). For the purpose of open access, the author has applied a CC-BY 4.0 public copyright license to all author accepted manuscripts (AAM) arising from this submission.

Authors thank Anushka Takhi, Joana Petushi, Laura Demmer and Zoe Fisk for assistance with the experiments and assays; Katharina Klinger for assistance with animal care and use requirements; personnel at the DZNE Light Microscope Facility, Image and Data Analysis Facility and Preclinical Center; and Sarah A. Jewell for her valuable comments on the manuscript.

## Supplementary Information

### Supplementary figure legends

**Supplementary Fig. 1.**
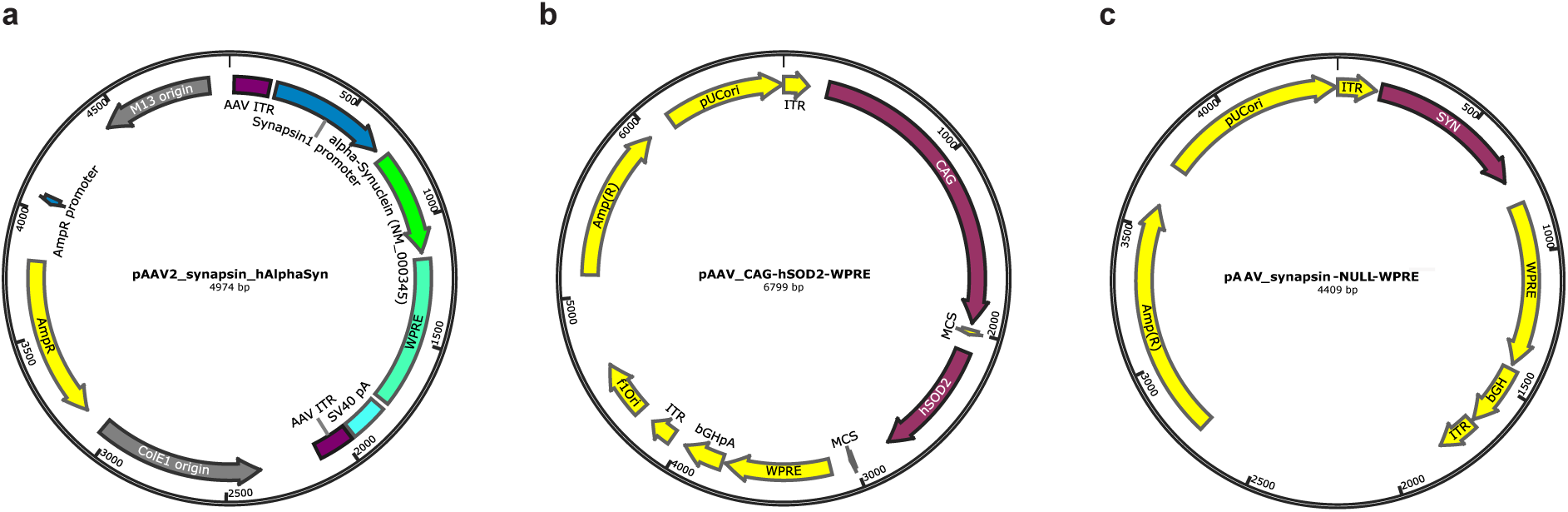
Maps of AAVs used in the study. Maps showing the expression cassettes of vectors used for transduction of hASYN (**a**) or SOD2 (**b**). The map of empty AAVs lacking protein coding sequence is also shown in (**c**).

**Supplementary Fig. 2.**
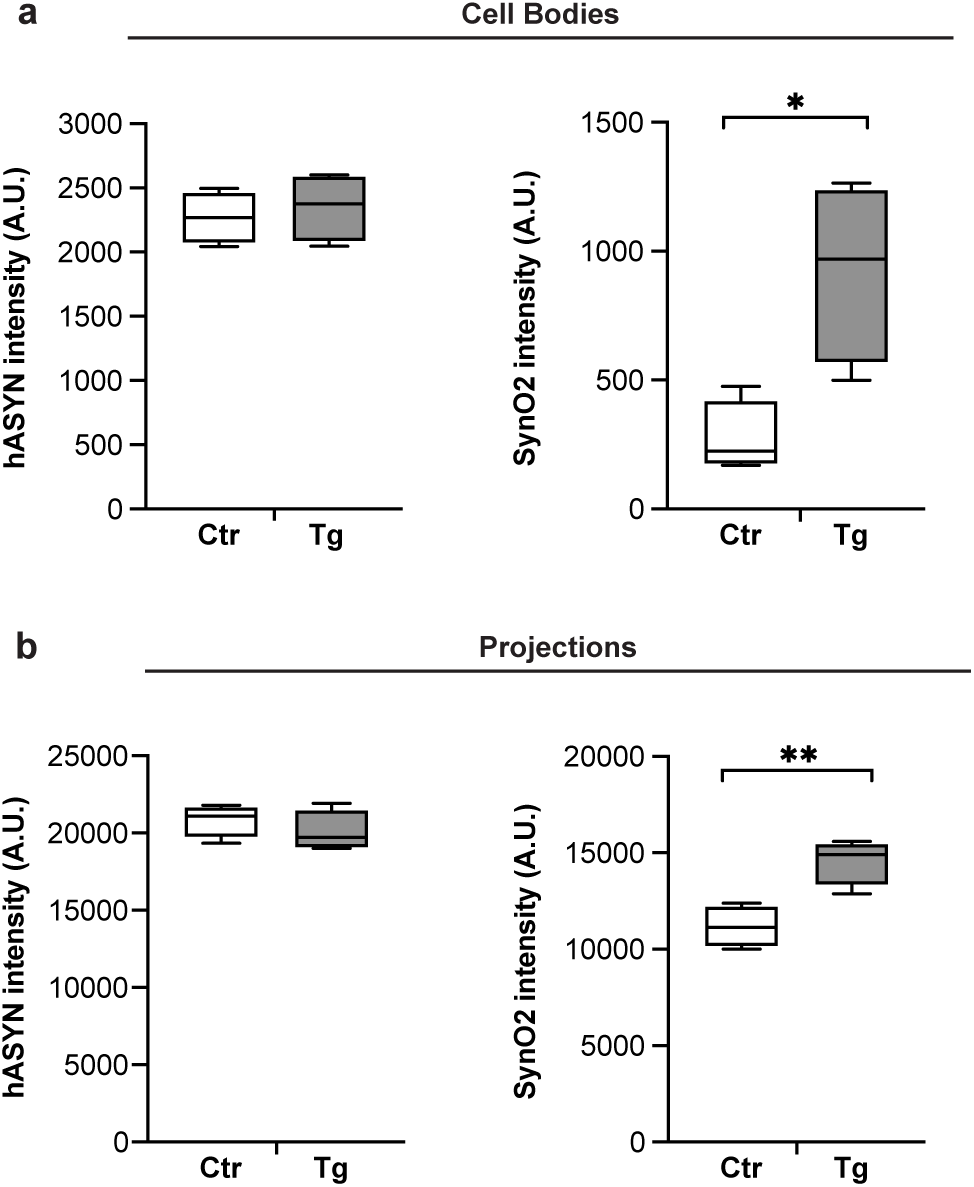
L444P-induced ASYN aggregation assessed by SynO2 staining. All mice received an intravagal injection of hASYN-delivering AAVs. MO tissue sections were double-stained with anti-hASYN and an antibody, SynO2, that reacts with aggregated ASYN species. **a, b** Fluorescent intensity (expressed as arbitrary units, A.U.) of hASYN and SynO2 signals were measured in hASYN-containing neuronal cell bodies (**a**) and projections (**b**) in the left DMnX of control (Ctr) and L444P mutant (Tg) mice (n = 4/group). Plots show median, upper and lower quartiles, and maximum and minimum as whiskers. \**P* :< 0.05 and \*\**P* :< 0.001, Student’s t-test.

**Supplementary Fig. 3.**
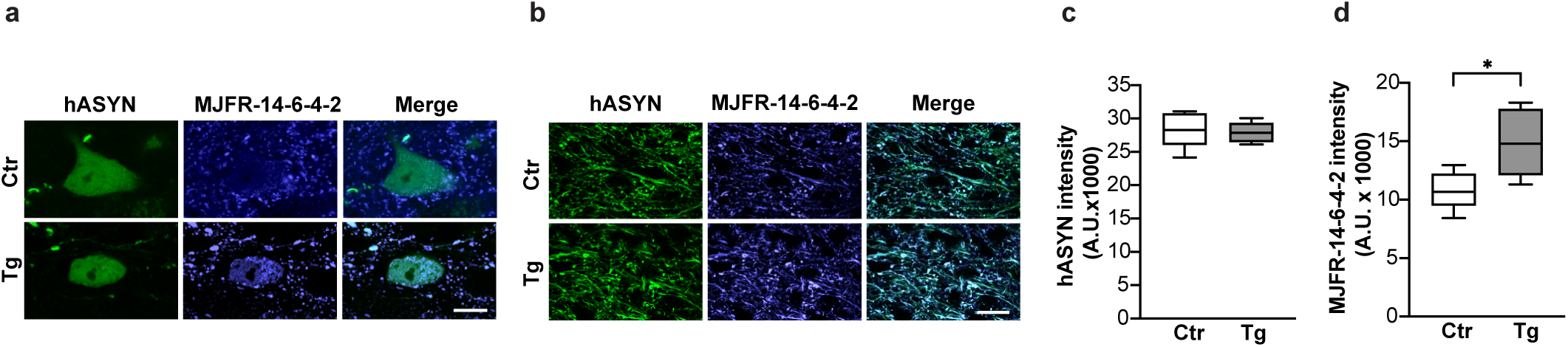
L444P-induced ASYN aggregation assessed by MJFR-14-6-4-2 staining. All mice received an intravagal injection of hASYN-delivering AAVs. MO tissue sections were double-stained with anti-hASYN and an antibody, MJFF-14-6-4-2, that reacts with aggregated ASYN species. **a, b** Representative fluorescent images of neuronal cell bodies (**a**) and projections (**b**) in the left (ipsilateral to the AAV injection) DMnX of a control (Ctr) and an L444P mutant (Tg) mouse. Scale bar: 10 μm (**a**) and 20 μm (**b**). **c, d** Fluorescent intensity (expressed as arbitrary units, A.U.) of hASYN immunoreactivity was measured in the left DMnX (**c**); fluorescent MJFF-14-6-4-2 intensity was measured within hASYN-containing DMnX neurons (both cell bodies and neurites) (**d**); samples were obtained from control and L444P mutant mice (n = 5/group). Plots show median, upper and lower quartiles, and maximum and minimum as whiskers. \**P* :< 0.05, Student’s t-test.

**Supplementary Fig. 4.**
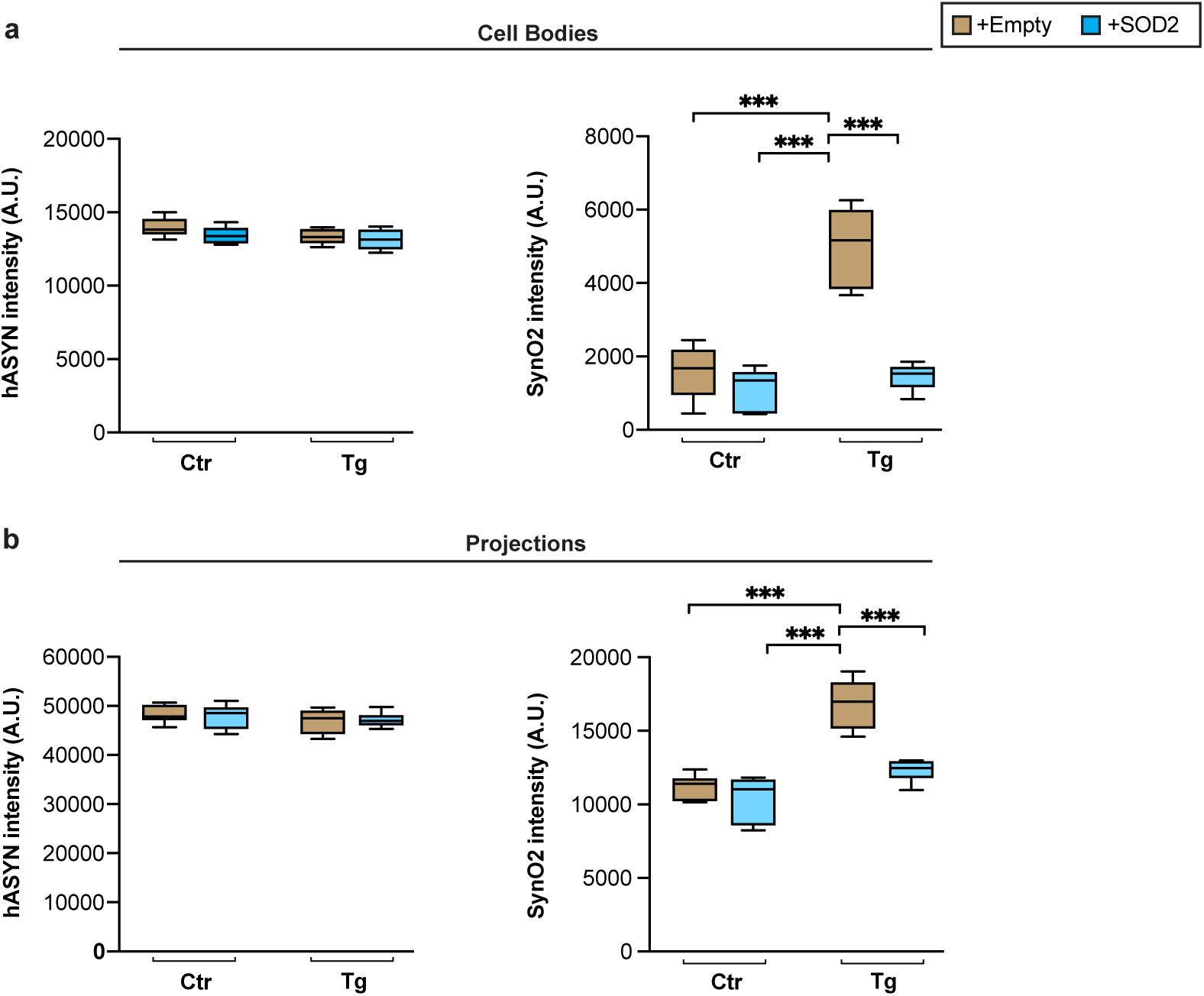
Effects of SOD2 transduction on L444P-induced ASYN aggregation. Control (Ctr) and L444P mutant (Tg) mice were all injected intravagally with hASYN-delivering AAVs. Groups of animals also received, together with hASYN-AAVs, empty vectors lacking protein coding (+empty) or AAVs delivering SOD2 DNA (+SOD2). MO tissue sections were double-stained with anti-hASYN and anti-SynO2. **a, b** Fluorescent intensity (expressed as arbitrary units, A.U.) of hASYN and SynO2 signal in hASYN-containing neuronal cell bodies (**a**) and projections (**b**) in the left DMnX of control and L444P mutant mice co-injected with either hASYN/empty (light brown)- or hASYN/SOD2(azure blue)-AAVs (n ≥ 5/group). Plots show median, upper and lower quartiles, and maximum and minimum as whiskers. * *P* :< 0.05 and \*\**P* :< 0.001, Tukey post-hoc test.

## References

1. Grabowski, G. A. Phenotype, diagnosis, and treatment of Gaucher’s disease. Lancet, 372, 1263–1271 (2008).

2. Goker-Alpan, O., Schiffmann, R., LaMarca, M. E., Nussbaum, R. L., McInerney-Leo, A., & Sidransky, E. Parkinsonism among Gaucher disease carriers. J Med Genet, 41, 937–940 (2004.)

3. Sidransky, E., et al. Multicenter analysis of glucocerebrosidase mutations in Parkinson’s disease. N Engl J Med, 361, 1651–1661 (2009).

4. Neumann, J., et al. Glucocerebrosidase mutations in clinical and pathologically proven Parkinson’s disease. Brain, 132, 1783–1794 (2009).

5. Vieira, S. R. L., & Schapira, A. H. V. Glucocerebrosidase mutations and Parkinson disease. J Neural Transm (Vienna), 129, 1105–1117 (2022).

6. Anheim, M., et al. Penetrance of Parkinson disease in glucocerebrosidase gene mutation carriers. Neurology, 78, 417–420 (2012).

7. Balestrino, R., Tunesi, S., Tesei, S., Lopiano, L., Zecchinelli, A. L., & Goldwurm, S. Penetrance of Glucocerebrosidase (GBA) Mutations in Parkinson’s Disease: A Kin Cohort Study. Mov Disord, 35, 2111–2114 (2020).

8. Nalls, M. A., et al. A multicenter study of glucocerebrosidase mutations in dementia with Lewy bodies. JAMA Neurol, 70, 727–735 (2013).

9. Spillantini, M. G., & Goedert, M. Neurodegeneration and the ordered assembly of alpha-synuclein. Cell Tissue Res, 373, 137–148 (2018).

10. Killinger, B. A., & Kordower, J. H. Spreading of alpha-synuclein - relevant or epiphenomenon? J Neurochem, 150, 605–611 (2019).

11. Mazzulli, J. R., et al. Gaucher disease glucocerebrosidase and alpha-synuclein form a bidirectional pathogenic loop in synucleinopathies. Cell, 146, 37–52 (2011).

12. Cullen, V., et al. Acid beta-glucosidase mutants linked to Gaucher disease, Parkinson disease, and Lewy body dementia alter alpha-synuclein processing. Ann Neurol, 69, 940–953 (2011).

13. Sardi, S. P., et al. Glucosylceramide synthase inhibition alleviates aberrations in synucleinopathy models. Proc Natl Acad Sci U S A, 114, 2699–2704 (2017).

14. Zunke, F., et al. Reversible Conformational Conversion of alpha-Synuclein into Toxic Assemblies by Glucosylceramide. Neuron, 97, 92–107 e110 (2018).

15. Rocha, E. M., et al. Glucocerebrosidase gene therapy prevents alpha-synucleinopathy of midbrain dopamine neurons. Neurobiol Dis, 82, 495–503 (2015).

16. Henderson, M. X., et al. Characterization of novel conformation-selective alpha-synuclein antibodies as potential immunotherapeutic agents for Parkinson’s disease. Neurobiol Dis, 136, 104712 (2020).

17. Migdalska-Richards, A., et al. The L444P Gba1 mutation enhances alpha-synuclein induced loss of nigral dopaminergic neurons in mice. Brain, 140, 2706–2721 (2017).

18. Yun, S. P., et al. alpha-Synuclein accumulation and GBA deficiency due to L444P GBA mutation contributes to MPTP-induced parkinsonism. Mol Neurodegener, 13, 1 10.1186/s13024-017-0233-5 (2018).

19. Li, H., et al. Mitochondrial dysfunction and mitophagy defect triggered by heterozygous GBA mutations. Autophagy, 15, 113–130 (2019).

20. Migdalska-Richards, A., et al. L444P Gba1 mutation increases formation and spread of alpha-synuclein deposits in mice injected with mouse alpha-synuclein pre-formed fibrils. PLoS One, 15, e0238075 (2020).

21. Johnson, M. E., et al. Heterozygous GBA D409V and ATP13a2 mutations do not exacerbate pathological alpha-synuclein spread in the prodromal preformed fibrils model in young mice. Neurobiol Dis, 159, 105513 (2021).

22. Polinski, N. K., et al. The GBA1 D409V mutation exacerbates synuclein pathology to differing extents in two alpha-synuclein models. Dis Model Mech, 15, (2022).

23. Mahoney-Crane, C. L., et al. Neuronopathic GBA1L444P mutation accelerates glucosylsphingosine levels and formation of hippocampal alpha-synuclein inclusions. J Neurosci, 43, 501–521 (2023).

24. Ulusoy, A., et al. Caudo-rostral brain spreading of alpha-synuclein through vagal connections. EMBO Mol Med, 5, 1119–1127 (2013).

25. Helwig, M., et al. Brain propagation of transduced alpha-synuclein involves non-fibrillar protein species and is enhanced in alpha-synuclein null mice. Brain, 139, 856–870 (2016).

26. Pinto-Costa, R., Harbachova, E., La Vitola, P., & Di Monte, D. A. Overexpression-induced alpha-synuclein brain spreading. Neurotherapeutics, 20, 83–96 (2023).

27. Helwig, M., et al. Neuronal hyperactivity-induced oxidant stress promotes in vivo alpha-synuclein brain spreading. Sci Adv, 8, eabn0356 (2022).

28. Musgrove, R. E., et al. Oxidative stress in vagal neurons promotes parkinsonian pathology and intercellular alpha-synuclein transfer. J Clin Invest, 129, 3738–3753 (2019).

29. Farfel-Becker, T., Do J., Tayebi N. & Sidransky E. Can *GBA1*-associated Parkinson disease be modeled in the mouse? Trends Neurosci. 42(9):631–43 (2019).

30. Vaikath, N. N., et al. Generation and characterization of novel conformation-specific monoclonal antibodies for alpha-synuclein pathology. Neurobiol Dis, 79, 81–99 (2015).

31. Martinez, T.N., et al. The Michael J. Fox Foundation’s strategy to generate, characterize, and distribute preclinical α-synuclein research tools for molecular biology. https://www.michaeljfox.org/publication/michael-j-fox-foundations-strategy-generate-characterize-and-distribute-preclinical (2016)

32. Kumar, S. T., Jagannath, S., Francois, C., Vanderstichele, H., Stoops, E., & Lashuel, H. A. How specific are the conformation-specific alpha-synuclein antibodies? Characterization and validation of 16 alpha-synuclein conformation-specific antibodies using well-characterized preparations of alpha-synuclein monomers, fibrils and oligomers with distinct structures and morphology. Neurobiol Dis, 146, 105086 (2020).

33. Roberts, R. F., Wade-Martins, R., & Alegre-Abarrategui, J. Direct visualization of alpha-synuclein oligomers reveals previously undetected pathology in Parkinson’s disease brain. Brain, 138, 1642–1657 (2015).

34. Cleeter, M. W., et al. Glucocerebrosidase inhibition causes mitochondrial dysfunction and free radical damage. Neurochem Int, 62, 1–7 (2013).

35. Braak, H., Del Tredici, K., Rub, U., de Vos, R. A., Jansen Steur, E. N., & Braak, E. Staging of brain pathology related to sporadic Parkinson’s disease. Neurobiol Aging, 24, 197–211 (2003).

36. Braak, H., Rub, U., Gai, W. P., & Del Tredici, K. Idiopathic Parkinson’s disease: possible routes by which vulnerable neuronal types may be subject to neuroinvasion by an unknown pathogen. J Neural Transm (Vienna), 110, 517–536 (2003).

37. Braak, H., de Vos, R. A., Bohl, J., & Del Tredici, K. Gastric alpha-synuclein immunoreactive inclusions in Meissner’s and Auerbach’s plexuses in cases staged for Parkinson’s disease-related brain pathology. Neurosci Lett, 396, 67–72 (2006).

38. Beach, T. G., et al. Multi-organ distribution of phosphorylated alpha-synuclein histopathology in subjects with Lewy body disorders. Acta Neuropathol, 119, 689–702 (2010).

39. Chandra, R., et al. Gut mucosal cells transfer alpha-synuclein to the vagus nerve. JCI Insight, 8, 10.1172/jci.insight.172192 (2023).

40. Manning-Bog, A. B., Schule, B., & Langston, J. W. Alpha-synuclein-glucocerebrosidase interactions in pharmacological Gaucher models: a biological link between Gaucher disease and parkinsonism. Neurotoxicology, 30, 1127–1132 (2009).

41. Gundner, A. L., et al. Path mediation analysis reveals GBA impacts Lewy body disease status by increasing alpha-synuclein levels. Neurobiol Dis, 121, 205–213 (2019).

42. Surmeier, D. J. Determinants of dopaminergic neuron loss in Parkinson’s disease. FEBS J, 285, 3657–3668 (2018).

43. Schildknecht, S., et al. Oxidative and nitrative alpha-synuclein modifications and proteostatic stress: implications for disease mechanisms and interventions in synucleinopathies. J Neurochem, 125, 491–511 (2013).

44. Giasson, B. I., et al. Oxidative damage linked to neurodegeneration by selective alpha-synuclein nitration in synucleinopathy lesions. Science, 290, 985–989 (2000).

45. Duda, J. E., et al. Widespread nitration of pathological inclusions in neurodegenerative synucleinopathies. Am J Pathol, 157, 1439–1445 (2000).

46. Paxinou, E., et al. Induction of alpha-synuclein aggregation by intracellular nitrative insult. J Neurosci, 21, 8053–8061 (2001).

47. Burai, R., Ait-Bouziad, N., Chiki, A., & Lashuel, H. A. Elucidating the Role of Site-Specific Nitration of alpha-Synuclein in the Pathogenesis of Parkinson’s Disease via Protein Semisynthesis and Mutagenesis. J Am Chem Soc, 137, 5041–5052 (2015).

48. Murray, J., Taylor, S. W., Zhang, B., Ghosh, S. S., & Capaldi, R. A. Oxidative damage to mitochondrial complex I due to peroxynitrite: identification of reactive tyrosines by mass spectrometry. J Biol Chem, 278, 37223–37230 (2003).

49. Davis, C. W., et al. Nitration of the mitochondrial complex I subunit NDUFB8 elicits RIP1- and RIP3-mediated necrosis. Free Radic Biol Med, 48, 306–317 (2010).

50. Baden, P., et al. Glucocerebrosidase is imported into mitochondria and preserves complex I integrity and energy metabolism. Nat Commun, 14, 1930 (2023).

